# Cereblon controls the timing of muscle differentiation in *Ciona* embryos

**DOI:** 10.1101/2022.08.16.504204

**Authors:** Juanjuan Long, Andrea Mariossi, Chen Cao, Michael Levine, Laurence A. Lemaire

## Abstract

Thalidomide has a dark history as a teratogen, but in recent years it has been shown to function as a chemotherapeutic agent. Thalidomide binds cereblon, a component of E3 ubiquitin ligase complex and modifies its degradation targets. Despite these insights, remarkably little is known about the normal function of cereblon in development. Here, we employ the simple proto-vertebrate model, *Ciona intestinalis*, to address this question. We observed a “hotspot” of *Crbn* expression in the developing tail muscles and identify its enhancer containing both Myod activator sites and a Snail repressive element. Overexpression of Crbn in tail muscles decreases expression of contractility genes. We suggest that this reduction is due to premature degradation of Tbx6. Drug inhibition studies using lenalidomide, a derivative of thalidomide, lead to a striking increase of *Crbn* expression. This autofeedback regulation could be induced by neomorphic degradation of Snail, contributing to the efficacy of lenalidomide in treating metastatic melanoma. In summary, our analysis of Crbn suggests that its normal function in *Ciona* is to time expression of contractility genes during muscle differentiation thereby ensuring coordination of tail morphogenesis and hatching of swimming tadpoles.

## INTRODUCTION

Cereblon (*CRBN*) is a component of an E3 ubiquitin ligase complex with a mottled history. It was implicated in severe birth defects triggered by the use of thalidomide during pregnancy (Ito et al., 2010; Lenz and Knapp, 1962; Mcbride, 1961). In particular, the human form of *CRBN*, but not the mouse or rat version, mediates the degradation of a spalt-like transcription factor 4 (SALL4) and promyelocytic leukemia zinc finger protein (PLFZ) upon association with thalidomide exerting teratogenic effects. Both are zinc finger transcription factors and important for the growth of the long bones of developing limbs, and their loss probably represents a major contributing factor to the birth defects caused by thalidomide in humans (Chen et al., 2020; Donovan et al., 2018; Matyskiela et al., 2018; Yamanaka et al., 2021). Thalidomide and its analogues such as lenalidomide have recently re-emerged as highly effective treatments against multiple myeloma and other B cell neoplasms due to their immunomodulatory activities (IMiDs) (Hideshima et al., 2000; Richardson et al., 2002). They are also used to treat neurodegenerative and neuropsychiatric disorders and hematopoietic malignancies (Bartlett et al., 2004; Muller et al., 1999). In myeloma cell lines, CRBN and lenalidomide (and derivatives) target degradation of the zinc transcription factors Ikaros (IKZF1) and Aiolos (IKZF3), causing arrest of cell proliferation and death (Gandhi et al., 2014; Krönke et al., 2014; Lu et al., 2014). Despite years of research aimed at understanding how cereblon influences these developmental and diseases processes little is known about its normal physiological activities.

Cereblon acts as a substrate receptor with the DNA damage-binding protein-1 (DDB1), cullin-4A and 4B (CUL4A and CUL4B), and the E3 ubiquitin ligase ring box 1 (RBX1) in the ubiquitinating complex (Angers et al., 2006; Ito et al., 2010). This complex appears to function in a highly pleiotropic manner to target protein degradation in vertebrate models such as zebrafish, chicken, and mice. In mouse embryos, *Crbn* expression is enriched in hippocampal and cerebellum regions of the developing brain as well as in serotoninergic and adrenergic neurons (Aizawa et al., 2011). *Crbn*-null mice do not show obvious developmental defects and are viable, although mutant mice have some cognitive defects akin to mild mental retardation anomalies in humans (Higgins et al., 2004; Rajadhyaksha et al., 2012). Mutant mice also display a surprising resilience to obesity and diabetes when raised on a high fat diet (Lee et al., 2013). In zebrafish, *crbn* is detected throughout embryogenesis and involved in the morphogenesis of the pectoral fin and otic vesicle (Ando et al., 2019; White et al., 2017). Knockdown of *crbn* leads to an enlarged head and aberrant eye development likely associated with lower Wnt signaling activity due to the ability of Cereblon to target Casein kinase 1a1, an essential component of the β catenin destruction complex (Ando et al., 2019; Shen et al., 2021). However, none of these studies uncovered the normal regulation and functions of cereblon during limb formation. Here, we employ the simple proto-vertebrate, *Ciona intestinalis*, to explore the normal activities of Crbn during muscle development.

*Ciona* embryo and larvae have been a favorite of experimental embryologists for over 150 years due to their small cell numbers, simple lineages and similarity of developmental programs with vertebrates (Conklin, 1905; Imai et al., 2006; Whittaker, 1977). Recent single cell RNA-sequencing (scRNA-seq) assays led to a comprehensive atlas of *Ciona* development, spanning the onset of gastrulation to the hatching of swimming tadpoles (Cao et al., 2019). We used these maps to identify the sites and timing of *Crbn* expression during development, and present evidence that it plays a role in controlling the timing of muscle differentiation in developing tadpoles. We also performed scRNA-seq on lenalidomide-treated embryos and provide evidence for feedback regulation of *Crbn* expression. Overall, we propose that Crbn controls the timing of muscle differentiation by modulating the degradation of key muscle transcription factors such as the T-box transcription factor Tbx6 and Snail.

## RESULTS

### Transient expression of *Crbn* in developing muscles

The *Ciona* genome contains a single orthologue of *Crbn*. As in humans, the *Ciona* thalidomide-binding domain contains a conserved glutamine 377 (E377) residue that is important for the interaction between CRBN and PLZF or SALL4 (Donovan et al., 2018; Matyskiela et al., 2018; Yamanaka et al., 2021). In humans, valine 388 (V388) is a key residue for stabilizing thalidomide-dependent CRBN interactions. However, the corresponding amino acid residue in *Ciona* is isoleucine, as seen in chicken, frog, and zebrafish (Supp Fig. 1A). These organisms display developmental defects upon treatment with lenalidomide and derivatives (IMiDs), including malformations of fins and limbs (Christian et al., 2007; Fort et al., 2000; Ito et al., 2010; Mahony et al., 2013; Matyskiela et al., 2018). Non responsive organisms like mouse and rat have two critical amino acid substitutions in this domain, E377V and V388I, that prevent stable associations with SALL4 and IKZF1/3 in the presence of IMiDs, and no limb defects are observed (Supp Fig. 1B) (Donovan et al., 2018; Matyskiela et al., 2018). In contrast to the ubiquitous profiles of *crbn* expression reported in zebrafish (Ando et al., 2019; White et al., 2017), we observed a hotspot of expression only in developing muscles in the *Ciona* scRNA-seq atlas (Fig. 1A and B) (Cao et al., 2019). Expression is first detected during neurula stages, reaches a peak in early and mid tailbud stages, and then diminishes in late tailbud embryos prior to the hatching of swimming tadpoles (Fig. 1C and D).

**Fig. 1:**
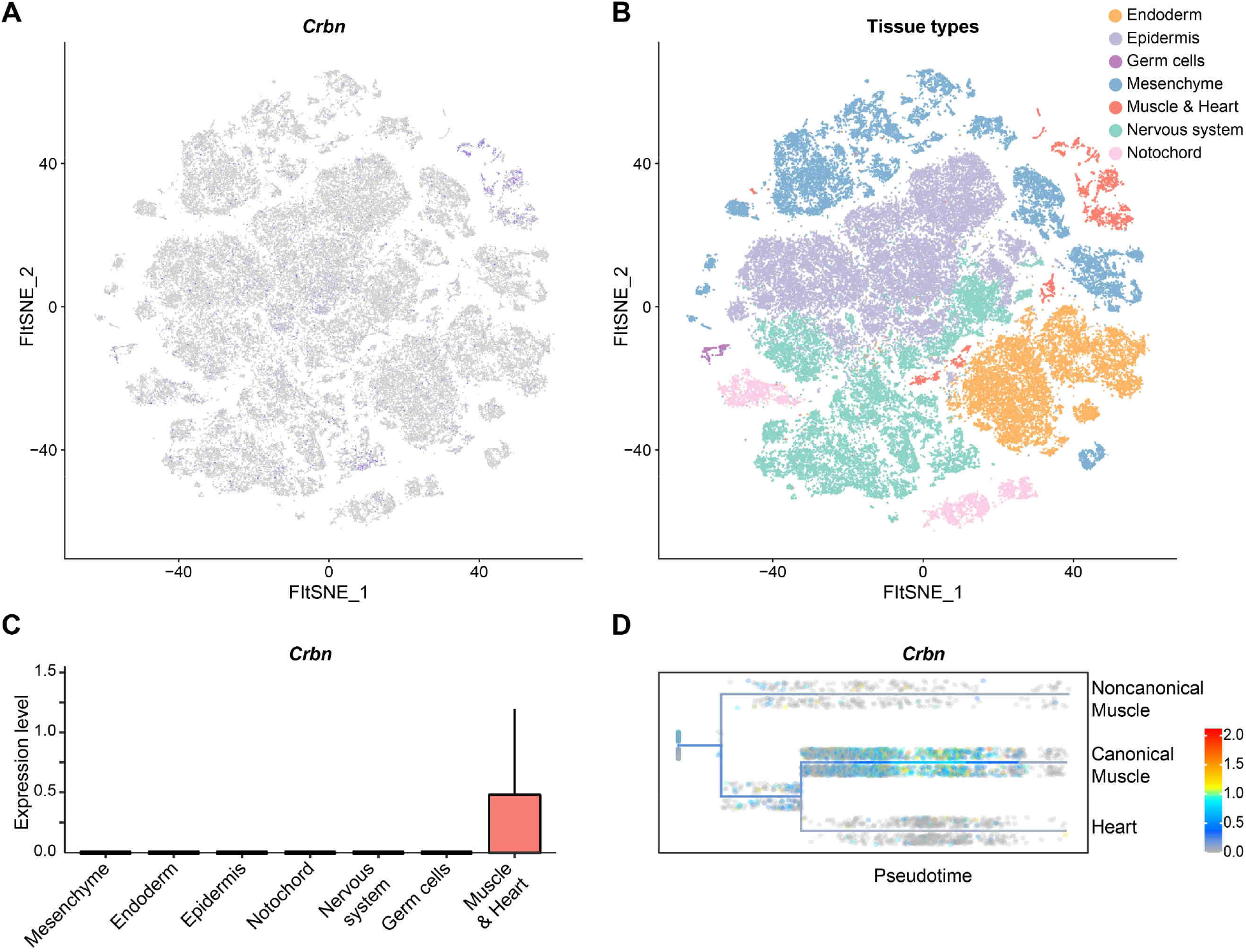
Expression of *Crbn* during *Ciona* embryogenesis. **A.** t-distributed stochastic neighbour-embedding (t-SNE) plot of *Crbn* expression in single cells across 10 different stages from the gastrula to the larva stage (n = 90 579 cells). **B.** t-SNE plot of single cells at the same stages, the cells are color coded by tissue types (n = 90 579 cells). **C.** Box plot of *Crbn* expression across tissue types (Mesenchyme: n = 19 143 cells; Endoderm: n = 14 162 cells; Epidermis: n = 26 936 cells; Notochord: n = 4 053 cells; Nervous system: n = 22 198 cells; Germ cells: n = 396 cells and Muscle & Heart: n = 3 691 cells). The color coding is the same as in B. **D.** Expression of *Crbn* along the muscle and heart lineages from the gastrula to the larva stage. The cells are ordered by pseudotime (n = 3 691 cells).

To verify the basis for this muscle-specific expression profile, we cloned a series of upstream regions of the initiating ATG codon into reporter genes expressing histone fused to green fluorescent protein (*H2BGFP*) (Fig. 2). We were able to identify a tissue-specific enhancer containing 455bp, 410bp, and 314bp of the 5’ flanking region, which direct robust expression in the tail muscles of electroporated embryos (Fig. 2B and C). These short sequences contain several putative regulatory elements, most notably, three myogenic determinant (Myod) binding motifs as well as a putative Snail repressor site (summarized in Fig. 2A). Similar to vertebrate muscle development, *Ciona* Myod specifies the muscle lineage (Meedel et al., 2007).

**Fig. 2:**
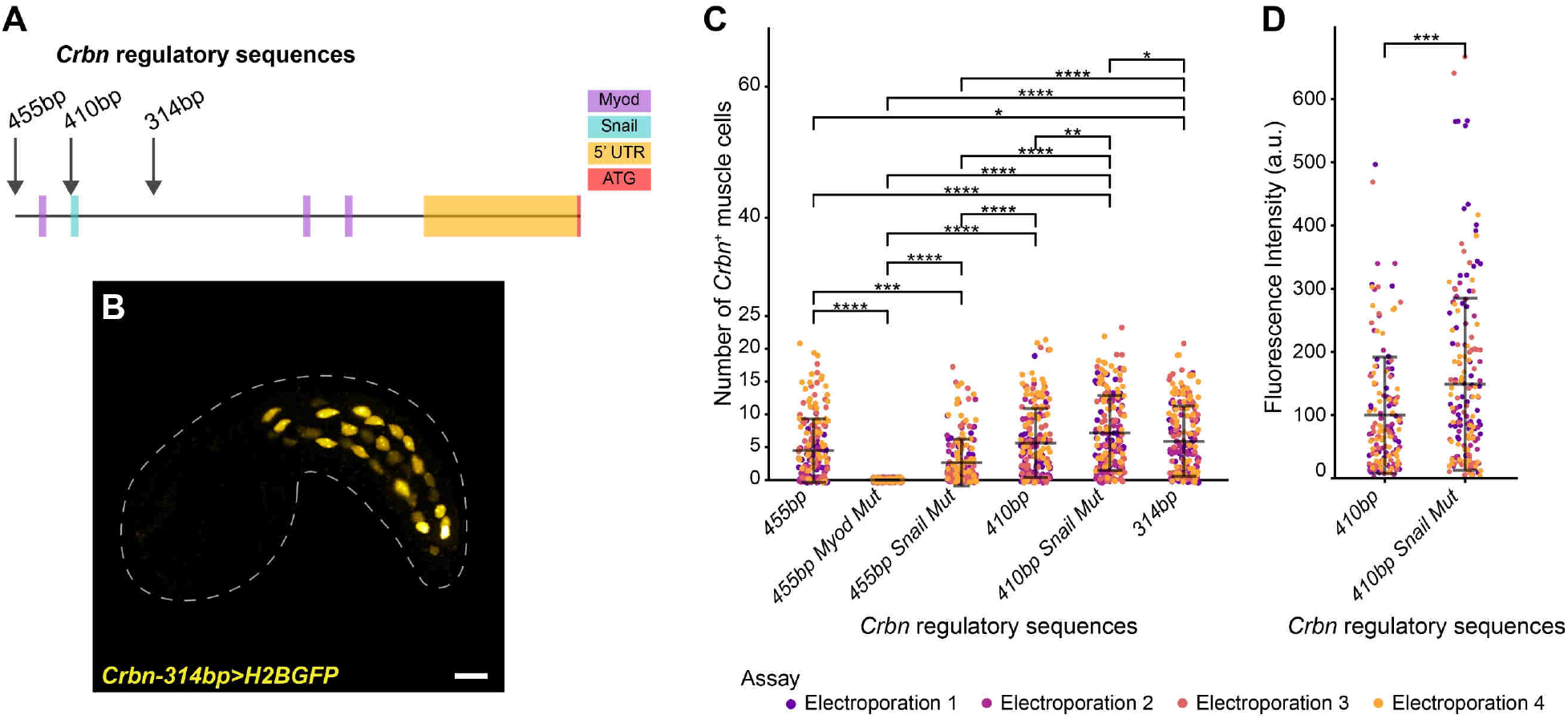
*Crbn* regulatory sequences. **A.** Diagram of *Crbn* regulatory sequences. The analyzed regulatory sequences were 455bp, 410bp and 314bp from the translation start site (red). The position of the binding motifs for Myod and Snail are indicated in purple and turquoise respectively, the 5’ untranslated region (5’UTR) is represented in orange. **B.** Representative embryo electroporated with the *Crbn* reporter genes consisting of the 314bp upstream sequence fused to histone GFP (H2BGFP). Scale bar: 20μm. **C.** Dot plot of the number of muscle cells expressing the different reporters. The embryos were pooled over 4 different electroporations. 455bp: 4.5 +/- 4.9, n = 206 embryos; 455bp Myod mut: 0 +/- 0, n = 208 embryos; 455bp Snail mut: 2.6 +/- 3.6, n = 208 embryos; 410bp: 5.6 +/- 5.3, n = 205 embryos; 410bp Snail mut: 7.2 +/- 5.7, n = 208 embryos; 314bp: 5.9 +/- 5.4 n = 207 embryos. Nested ANOVA with random effect (Assay) with 5 degrees of freedom followed by pairwise comparison with Tukey adjustment (degree of freedom 1235) (p value in Supp Table 1). **D.** Dot plot of the total fluorescence intensity for 410bp enhancer reporter (100 +/- 92, n = 205 embryos) and 410bp Snail mut enhancer reporter (149 +/- 136, n = 208 embryos). Nested ANOVA with random effect (Assay) with 1 degrees of freedom: p value = 1×10^-4^; random effect: p value = 0.1. All data are shown as mean +/- standard deviation; *, p value < 5×10^-2^; **, p value < 1×10^-2^; ***, p value < 1×10^-3^; **** p value < 1×10^-4^.

To test if these binding sites contributed to the expression of *Crbn*, we systematically mutagenized them in reporter genes and visualized their activities either by quantifying the number of muscle cells expressing the reporter or measuring total fluorescence intensity (Fig. 2C and D, Supp Table 1). Most notably, mutations in all three Myod sites abolished the expression of the 455bp reporter gene (Fig. 2C, Supp Table 1), while mutations in the lone Snail site resulted in a slight increase in the fluorescence intensity of the 410bp reporter (Fig. 2D). These results are consistent with the hypothesis that Myod and Snail directly activates and represses, respectively, the *Crbn* enhancer to produce a transient expression profile during muscle development.

### Muscle differentiation is delayed upon *Crbn* overexpression

To assess the role of *Crbn* in developing tail muscles, we focused on overexpression assays since thalidomide and its derivatives cause Crbn to recruit neomorphic substrates (Gandhi et al., 2014; Krönke et al., 2014; Lu et al., 2014). Towards this end, we examined the consequences of targeted overexpression of the endogenous *Ciona Crbn* gene in developing tail muscles using the muscle-specific enhancer of *Tbx6b* (Takatori et al., 2004). *Tbx6b* expression starts in the presumptive muscles before the onset of gastrulation. Overexpression of *Crbn* in developing muscles does not cause any obvious morphological defects in larvae (Supp Fig. 2), although it is possible that the shapes of the individual muscle cells are slightly less rectangular in transgenic embryos at earlier stages evocative of less mature cells (Fig. 3A and B).

**Fig. 3:**
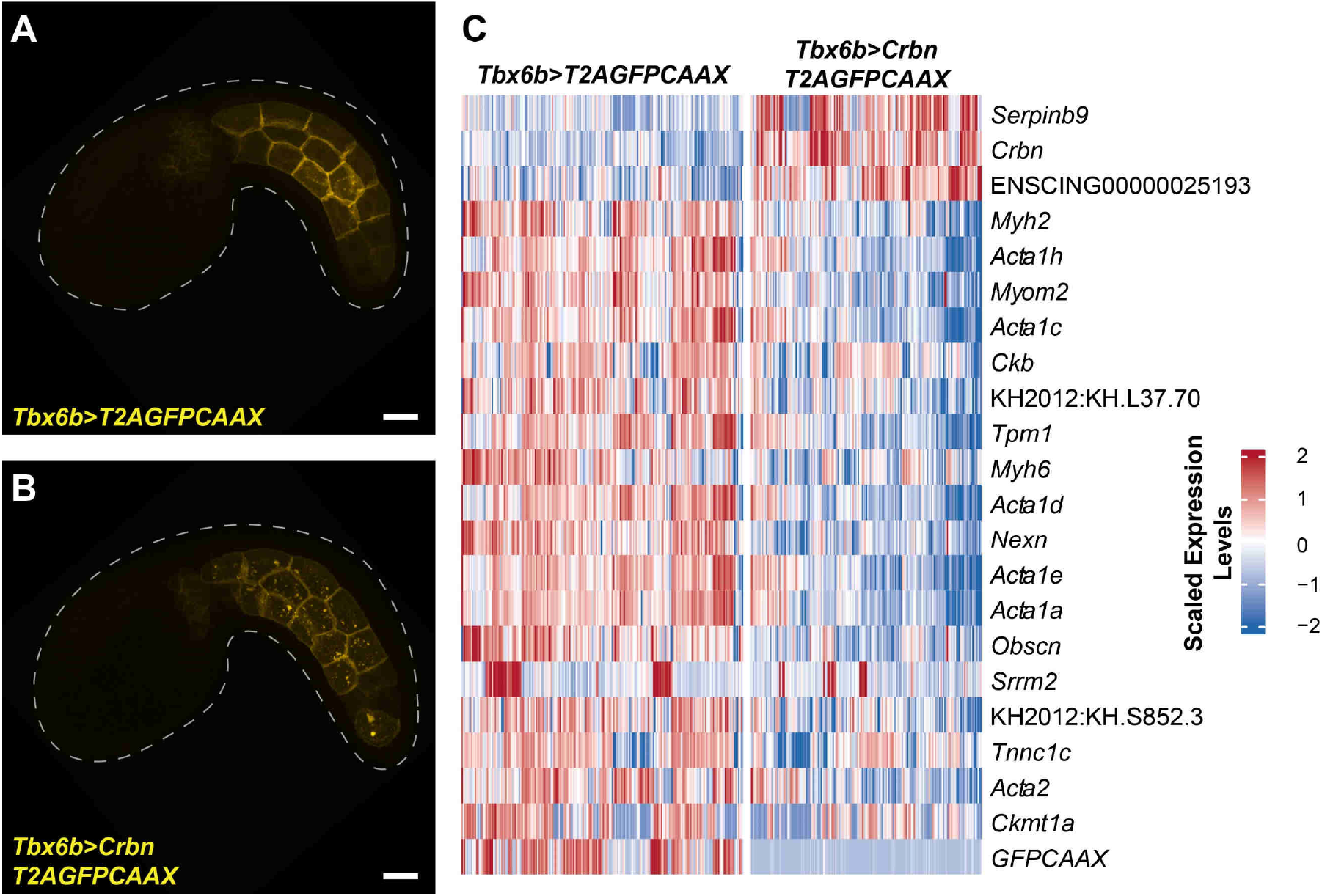
*Crbn* overexpression in *Ciona* muscle cells. **A.** and **B.** Representative mid tailbud embryos expressing either *GFP* targeted to the membrane (*T2A*GFPCAAX) (A.) or *Crbn* and membrane *GFP (CrbnT2AGFPCAAX*) (B.) under the muscle specific regulatory sequences of *Tb6xb*. **C.** Heatmap of the genes differentially expressed with at least 1.4-fold change and an adjusted p value below 0.05 between *Ciona* muscle cells expressing the transgene *Tbx6b>T2AGFPCAAX* (control) and the transgene *Tbx6b>CrbnT2AGFPCAAX (Crbn* overexpression). Scale bar 20 μm.

To investigate the consequences of *Crbn* overexpression in early muscle differentiation, we took advantage of one of the key attributes of the *Ciona* system, namely, its small cells number and use of scRNA-seq as a primary screening tool. Electroporated embryos expressing either *Tbx6b>Crbn* (*Crbn* overexpressing embryo) or *Tbx6b>GFP* (control embryos) were raised to mid tailbud stages, and individual cells were dissociated and sequenced to ^~^12.3-fold coverage, permitting identification of all cell types. After data processing and cell type assignment using known markers (Supp Fig. 3A to D) (Cao et al., 2019; Imai et al., 2006), we could verify that overexpression of *Crbn* did not prevent specification of muscle cells since their numbers were not affected (Supp Fig. 3E). However, the transcriptome profiles of *Crbn* overexpressing embryos displayed a clear deviation from control embryos (Supp Table2, Supp Fig. 3F). There is a significant reduction in the expression of a variety of muscle structural genes such as troponin C1 (*Tnnc1c*), different actin alpha 1 genes (*Acta1a*, *Acta1c*, *Acta1d*, *Acta1e*, *Acta1h*), and tropomyosin 1 (*Tpm1*) (Fig. 3C). These results are consistent with the possibility that *Crbn* leads to a delay in muscle differentiation by attenuating the expression of muscle effector genes. Hatching of swimming tadpoles depends on coordinated contractions of the tail muscles, and *Crbn* might prevent these contractions from occurring prematurely during tailbud stages. Most of these genes are activated by Tbx6 and Myod (Yu et al., 2019), which could be a target of Crbn-mediated proteasomal degradation (see Discussion).

### *Crbn* is upregulated upon lenalidomide treatment

Lenalidomide was designed to be more potent and with potentially fewer side effects (Bartlett et al., 2004; Richardson et al., 2002). Like thalidomide, its primary target is Crbn (Gandhi et al., 2014; Krönke et al., 2014; Lu et al., 2014). To gain further insights into its effects and putative mechanisms of action, we treated embryos with lenalidomide and N-methyl lenalidomide, a negative control since the additional N-methyl group prevents binding to Cereblon (Akuffo et al., 2018). Treated late tailbud embryos exhibited upward bending of the tails, contrary to the downward bending observed in normal embryos. This phenotype is consistent with subtle changes in the morphology of individual muscle cells as observed in *Crbn* overexpressing experiments (Fig. 4A to C). Increasing amounts of each drug led to a progressive inhibition of tail muscle contractions in swimming tadpoles (Fig. 4D and E). The concentration at which 50% of the larvae fail to twitch their tails (IC50) was 291 and 316 μM for lenalidomide and N-methyl lenalidomide, respectively. These concentrations are on the same order as those found for zebrafish (Mahony et al., 2013).

**Fig. 4:**
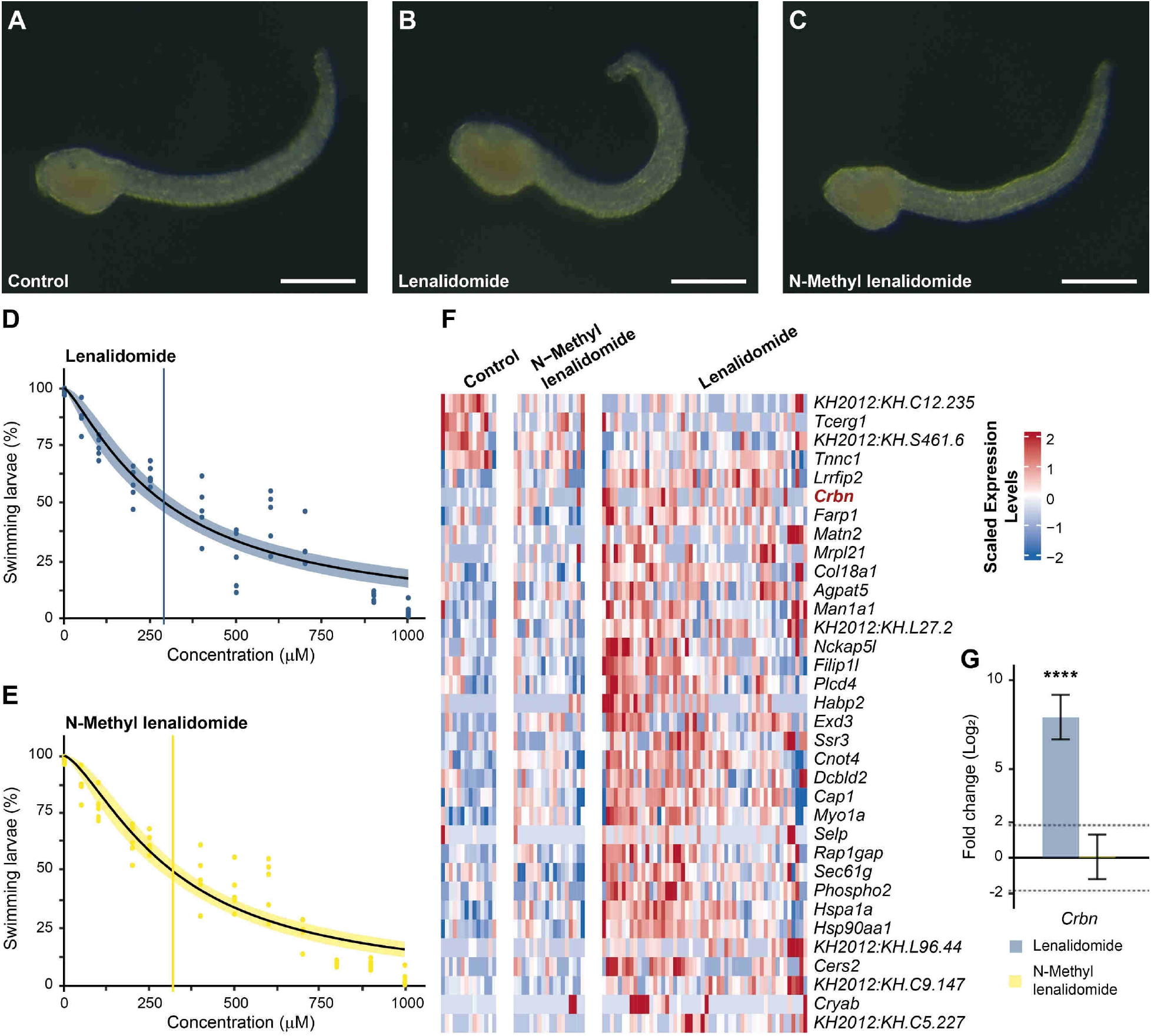
Effect of lenalidomide and N-Methyl Lenalidomide on muscle development. **A.** to **C.** Examples of late tailbud 2 embryos treated with DMSO (A.), 200 μM of lenalidomide (B.), or 200 μM of N-Methyl lenalidomide (C.) since the gastrula stage. Lenalidomide exposure resulted in an upward tail and/or abnormal tip development. **D.** and **E.** Dose - response curve of lenalidomide (D.) and N-Methyl lenalidomide (E.) showing the percentage of larvae twitching their tail in a 30s window in function of the concentration of the drug. The vertical bars correspond to the IC50 for lenalidomide (291 μM, D.) and N-Methyl lenalidomide (316 μM, E.) respectively (n = 5). **F.** Heatmap of the 30 most differentially expressed genes (up and down) with at least 1.4-fold change (p value below 0.05) in the muscle cells of late tailbud 2 embryos treated with 200 μM of lenalidomide compared to DMSO and 200 μM of N-Methyl lenalidomide treated embryos. *Crbn* is highlighted in red. **G.** qPCR analysis for *Crbn* on late tailbud 2 embryos treated with either 200 μM of lenalidomide or 200 μM of N-Methyl lenalidomide compared to DMSO treated embryos. (Mean +/- 95% CI: Lenalidomide p value < 2×10^-15^, n = 5; N-Methyl lenalidomide p value = 0.99, n = 5). Scale bar 50μm

To determine the basis for muscle defects, we raised embryos from the gastrula stage in 200 μM lenalidomide, N-methyl lenalidomide or DMSO control and conducted scRNA-seq assays. At the late tailbud stage, prior to hatching, embryos were dissociated and prepared for single-cell sequencing. Altogether, we obtained 1.4, 0.5, and 0.4-fold coverage for muscle cells with the three treatments (Supp Fig. 4). Several genes appear to be upregulated upon treatment with lenalidomide (Fig. 4F, Supp Fig. 5A, Supp Table 3 and 4). Most notably, *Crbn* itself exhibits upregulation upon addition of lenalidomide but not N-methyl lenalidomide. We confirmed this result by qPCR assays. *Crbn* displays a 256-fold increase as compared with control embryos (Fig. 4G). Moreover, cullin-5 (*Cul5*), another component of the E3 ubiquitination complex was also increased upon exposure to lenalidomide suggesting a broader effect on the pathway (Supp Fig. 5B). As discussed earlier, the muscle-specific enhancer regulating *Crbn* expression appears to be activated by Myod and repressed by Snail. It is possible that lenalidomide triggers the selective degradation of Snail by the Crbn E3 complex, leading to the observed increase in Crbn expression (summarized in Fig. 5).

**Fig. 5:**
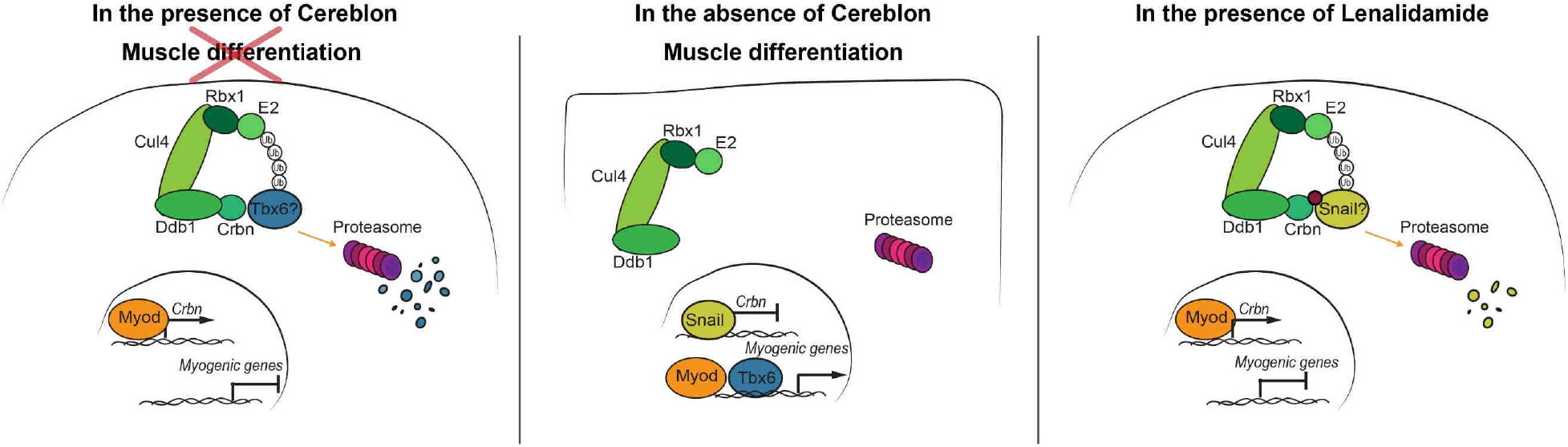
Role of *Crbn* in *Ciona* muscle cells. Myod activates *Crbn* gene expression. Crbn then as part of the E3 Ubiquitin ligase complex targets Tbx6 for degradation preventing muscle structural genes expression. When, Snail is repressing *Crbn* expression, Tbx6 then can activate muscle functional genes. In presence of lenalidomide, *Crbn* is overexpressed. It could be due to the Lenalidomide-Crbn containing E3 Ubiquitin ligase complex triggering the degradation of Snail, the repressor of *Crbn* leading to its upregulation.

## DISCUSSION

Cereblon is a highly conserved E3 ligase substrate receptor implicated in learning and memory, and associated with multiple roles in human health and disease (Kim et al., 2016). Here, we presented evidence that Crbn is specifically expressed in developing tail muscles of *Ciona* embryos. This expression is mediated by a tissue-specific enhancer located upstream of the *Crbn* coding sequence. The *Crbn* muscle enhancer contains binding sites for Myod and Snail, and targeted mutations in these sites cause changes in the activity of the enhancer as shown with reporter assays. scRNA-sequencing assays documented a striking attenuation in the transcript levels for a number of muscle structural genes upon targeted overexpression of *Crbn*. Drug inhibition studies raise the possibility of an autoregulatory feedback loop since treatment with lenalidomide leads to increased *Crbn* expression in late tailbud embryos. This effect is likely via Crbn since it is not observed when the embryos are exposed to N-methyl lenalidomide, the control drug unable to bind to Crbn (Akuffo et al., 2018). Overall, these studies suggest that a key function of the Crbn E3 ubiquitin complex is to delay muscle differentiation to achieve correct timing of the muscle contractions that promote hatching of swimming tadpoles. It is conceivable, but speculative, that altering Crbn function might result in heterochrony, whereby the different tissues forming the limb do not develop synchronously. Such asynchrony could contribute to the birth defects caused by thalidomide in humans.

The classical B4.1 lineage forms tail muscles due to localized inheritance of the maternal Macho-1 transcription factor, which in turn activates Tbx6 at the 32-cell stage to initiate muscle differentiation (Kugler et al., 2010). Tbx6 is encoded by four different genes namely *Tbx6a*, *Tbx6b*, *Tbx6c* and *Tbx6d*. All of these genes are transiently transcribed during the early phases of muscle development but give way to myogenic regulatory factors such as Myod at the onset of muscle differentiation. Indeed, Tbx6 activates Myod and together they activate muscle structural genes such as troponin (Yu et al., 2019). However, Tbx6 proteins are quickly degraded, and thereafter, Myod and Tbx15/18/22 are responsible for maintaining the expression of muscle structural genes (Yu et al., 2022). *Crbn* overexpression leads to the downregulation of these genes at mid tailbud stages. An attractive target for Crbn would be Myod forming a negative feedback loop with the latter. However, Myod protein levels are relatively stable during tailbud stages (Yu et al., 2022). Therefore, we propose that the Crbn E3 complex normally targets Tbx6 for degradation to reinforce the transition from muscle specification to differentiation. The resulting reduction in Tbx6 might also contribute to the delay in the onset of muscle contractility (Fig. 5). Future studies could explore the exact dynamics of Crbn protein synthesis, enzymatic activities and protein stability as well as its putative regulation of Tbx6. We predict a transient profile of activity extending from neurulation to mid tailbud development. Afterwards, *Crbn* levels diminish in parallel with Tbx6 in order to achieve full induction of muscle structural genes by Myod and Tbx15/18/22 prior to hatching.

It is possible that the normal reduction in Crbn expression and activity during late tailbud stages is due to the Snail repressor. Mutagenesis of the Snail binding site in the 410bp *Crbn* enhancer augmented expression of a GFP reporter gene. Moreover, addition of lenalidomide led to an increase in the levels of *Crbn* transcripts in late tailbud embryos (summarized in Fig. 5). This upregulation was absent when the embryos were treated with N-methyl lenalidomide suggesting that the effect is via Crbn itself (Akuffo et al., 2018). This increase could be due to the targeting of the Snail repressor by the Cereblon-lenalidomide complex. Interestingly, Snail is also a zinc finger transcription factor like SALL4, IKZF1, IKZF3 and PLZF. It is possible that thalidomide and its analogues triggers a neomorphic activity by linking Crbn to one of the Snail zinc fingers (Donovan et al., 2018; Matyskiela et al., 2018; Sievers et al., 2018; Yamanaka et al., 2021). Snail has been implicated as a major driver of tumorigenesis by promoting epithelial-mesenchyme transitions (EMT) and therefore metastasis (Wang et al., 2013). Further studies will determine whether degradation of Snail by lenalidomide contributes to its anti-cancer effects in metastatic melanoma.

## MATERIALS

### Ciona handling, embryo collection and electroporation

Adult *Ciona intestinalis* type A (Pacific species, also called *Ciona robusta*) were purchased from M-Rep, San diego, California. Gamete collection and fertilization were done as described in (Christiaen et al., 2009b). Between 40 and 100 μg of plasmid DNA were electroporated into 1-cell stage embryos as describe in (Christiaen et al., 2009a). The embryos were then raised until the desired stage before being fixed with MEM-FA (4% formaldehyde, 0.1 M Mops (pH 7.4), 0.5 M NaCl, 1 mM EGTA, 2 mM MgSO4, and 0.05% Tween-20) for 30 minutes at room temperature, washed several times in phosphate-buffered saline with 0.01% Tween-20, and mounted on slides using FluorSave Reagent (Millipore). To visualize the myofibers, actin was stained using ActinRed (ThermoFisher) before mounting the embryos. The electroporations were performed in at least in triplicates.

### Molecular Cloning

The KH and KY models of the genes mentioned in this manuscript are indicated in Supp Table 5 along with common synonyms and the closest human gene. The regulatory sequences of *Crbn* were PCR amplified from genomic DNA (primers in Supp Table 5) and cloned into a pCESA vector upstream of *H2BGFP* coding sequences using the restriction enzymes AscI and NotI (NEB England). Mutations into the Snail binding motif of the *Crbn* regulatory sequences were obtained by recombination using NEBuilder (NEB England) of PCR amplified fragments of the *Crbn* reporter constructs (primers in Supp Table 6). A gblock (IDT, sequence in Supp Table 7) of the *Crbn* 455bp enhancer containing the mutated Myod binding sites was recombined into a pCESA vector upstream of *H2BGFP* coding sequence. *Tbx6b* regulatory sequences were PCR amplified from genomic DNA and cloned into a pCESa vector by ligation using AscI and NotI upstream of the coding sequence of mouse *Cd4* fused to *GFP* (*CD4GFP*) (Christiaen et al, 2008) or AscI and EcoRI (NEBEngland) upstream of the coding sequence of the self-cleaving peptide *T2A* (Szymczak and Vignali, 2005) followed by *GFP* and the palmitoylation motif *CAAX* (*T2AGFPCAAX*) (primers in Supp Table 6). The *T2AGFPCAAX* coding construct was obtained by recombining the *Ciona Rcl1* coding sequence amplified from *Ciona* embryonic cDNA and the coding sequence for *T2AGFPCAAX* amplified from a *GFPCAAX* construct (Wagner and Levine, 2012) into a pCesA vector downstream of the regulatory sequences of *Msx* (Russo et al., 2004) (primers in Supp Table 6). The coding sequence of *Crbn* was amplified from *Ciona* embryonic cDNA (primers in Supp Table 6) and cloned into a pCESA plasmid downstream of *Tbx6b* regulatory sequences using the restriction enzymes NotI and FseI (NEBEngland) or NotI and EcoRI when *Crbn* was followed by the *T2AGFPCAAX* coding sequence.

### Drug treatment

To calculate the lowest possible dose of drugs that have a toxicological impact on *Ciona* development a 100 μM stock solution of lenalidomide (ThermoFisher) and N-methyl lenalidomide (Bristol Myers Squibb) was prepared in DMSO and further diluted to prepare 10 working concentrations from 10 nM to 1000 nM. Fertilized eggs were placed in 6-well plates (BD Falcon, Corning) with 40 embryos per well/group in 4 mL of filtered sea water and kept in an incubator at 18°C. At gastrulation, the stock solution of the drugs was separately added to each well. Control wells were treated with only DMSO. Development and locomotion from mid tailbud to larvae were assessed under a Nikon SMZ1500 stereomicroscope. We categorized whether a larva moved in a window of 30 seconds and assigned a binary code (binomial response) of 0 (no movement) and 1 (detection of active locomoting). Data from 5 independent measurements (embryos from different adult pairs), each calculated as an average of 15 larvae replicates per well per experiment, were collected and used to evaluate dose response curves. Slope, IC50 and 95% confidence interval (CI) were estimated for each compound using a three-parameter log-logistic function of the R package drc version 3.0-1 (Ritz et al., 2015; Team, 2015). compParm() function in drc was used to identify statistically significant difference in the IC50 values of the drug.

### Image acquisition and analysis

Drug treated embryos were imaged using a Leica M165SC stereomicroscope mounted with a Plan APO 1.0X M camera. Pictures for the reporter assays were acquired using a Zeiss 880 confocal microscope. The larval myofibers were imaged at high magnification with the Airyscan module of the Zeiss 880 and the images were processed with the default setting. Quantification of the number of GFP^+^ nuclei and the total fluorescence intensity were done using custom scripts in Fiji (ImageJ) (Schindelin et al., 2012). For the reporter assays, the embryos from different electroporations were pooled and the difference between the enhancers was tested statistically by a nested ANOVA followed by post hoc tests with the electroporation as nested factor using the R packages lme4 version 1.1-30 (Bates et al., 2015), lmerTest version 3.1-3 (Kuznetsova et al., 2017), nlme version 3.1-157 (Pinheiro et al., 2022; Pinheiro and Bates, 2000) and lsmeans version 2.30-0 (Lenth, 2016).

### scRNA-seq and analysis

1-cell stage embryo were electroporated either with *Tbx6d>T2AGFPCAAX* or *Tbx6d>CrbnT2AGFPCAAX*. GFP^+^ embryos were selected and dissociated at the mid tailbud stage as previously described in (Cao et al., 2019). For the drug treated single cell experiment, 200 μM of lenalidomide or N-Methyl lenalidomide was added to embryos at the gastrula stage, control embryos were exposed to the same amount of DMSO as the drug-treated embryos. The embryos were then raised until the late tailbud 2 stage before being dissociated as above. In both single cell experiments, between 1000 and 2000 cells were loaded onto the Chromium Controller (10x Genomics). The libraries were prepared as described in (Lemaire et al., 2021) and sequenced on Illumina NovaSeq 6000 with the SP reagent kit (100 cycles, paired-end). Filtering, quality control of the raw sequencing and generation of the FASTQ files were performed as described in (Cao et al., 2019). To generate the gene barcode matrices, the count pipeline was run on 10x Cell Ranger version 3.0.0 for the *Crbn* overexpressing scRNA-seq experiment and version 6.0.1 for drug treated experiment with the *Ciona intestinalis* type A reference sequences obtained from the Ghost database (KH model 2012) (Satou et al., 2005).

For each single cell experiments, the data were integrated using the RPCA method and further analyzed with Seurat version 4.1 (Butler et al., 2018). Once the muscle cells were identified, Wilcox tests with the default settings were performed to identify the differentially expressed genes between the conditions using Seurat. The heatmaps were plotted with the R package ComplexHeatmap version 2.10.0 (Gu et al., 2016).

*Ciona Crbn* expression pattern across 10 different developmental stages as well as its expression along the muscle lineage pseudotime was obtained from the expression matrices and analysis published in (Cao et al., 2019).

### RNA isolation, cDNA synthesis and qPCR

Fertilized embryos were raised at 18°C to late tailbud 2 stage as described above in 6-well plates with artificial seawater containing either DMSO, lenalidomide (200 μM) or N-methyl lenalidomide (200 μM). At the time of sample collection, 50 embryos were placed in 1.5ml Eppendorf tubes with 600 μl of TRIzol reagent (Invitrogen). Total RNA was isolated with Direct-zol RNA MiniPrep kit (Zymo) with 15 min DNase I digestion step according to the manufacturer’s instructions. A total amount of 0.5 μg of RNA was used for cDNA synthesis, employing iScript cDNA Synthesis Kit (Bio-rad). qPCR was performed on an ViiA 7 (Applied Biosystems) with SYBR green fluorescent label (Thermo Fisher). Analyses were performed with the R package MCMC.qpcr version 1.2.4 (Matz et al., 2013) which employs a Bayesian approach to data analysis. A model was fitted assuming no variation in the control gene *Eef1a* (Sekiguchi et al., 2020) and results are reported as log(2) fold-changes to DMSO state with Bayesian 95% credible intervals. Results are from three or four biological replicates. The primer sequences are listed in Supp Table 6.

### Data availability

Data presented in this paper are available upon request. The raw sequencing data and gene expression matrices will be available on GEO upon publication.

## ACKNOWLEDGMENTS

We thank members of the Levine laboratory for helpful discussions and feedback on the manuscript. We thank the Lewis-Sigler Institute Genomics Core Facility as well as the Bioinformatics Office for technical support. The authors would like also to acknowledge Joel W Thompson and Zhongying Mo from Bristol Myers Squibb for insightful discussions and sharing reagents. Cartoons in Supp Fig. 1B were created using BioRender.com. This study was funded by a National Institute of Health (NIH) grant to M.L. (NS076542) and by a grant from Celgene (Bristol Myers Squibb).

## AUTHOR CONTRIBUTION

JL and ML conceived the project. JL, AM, ML, LAL designed the experiments. JL, AM and LAL performed the experiments. CC and LAL performed the computational data analysis. All authors contributed to the interpretation of the results. AM, LAL and ML wrote the manuscript.

## SUPPLEMENTARY FIGURES AND LEGENDS

**Supp Fig. 1:**
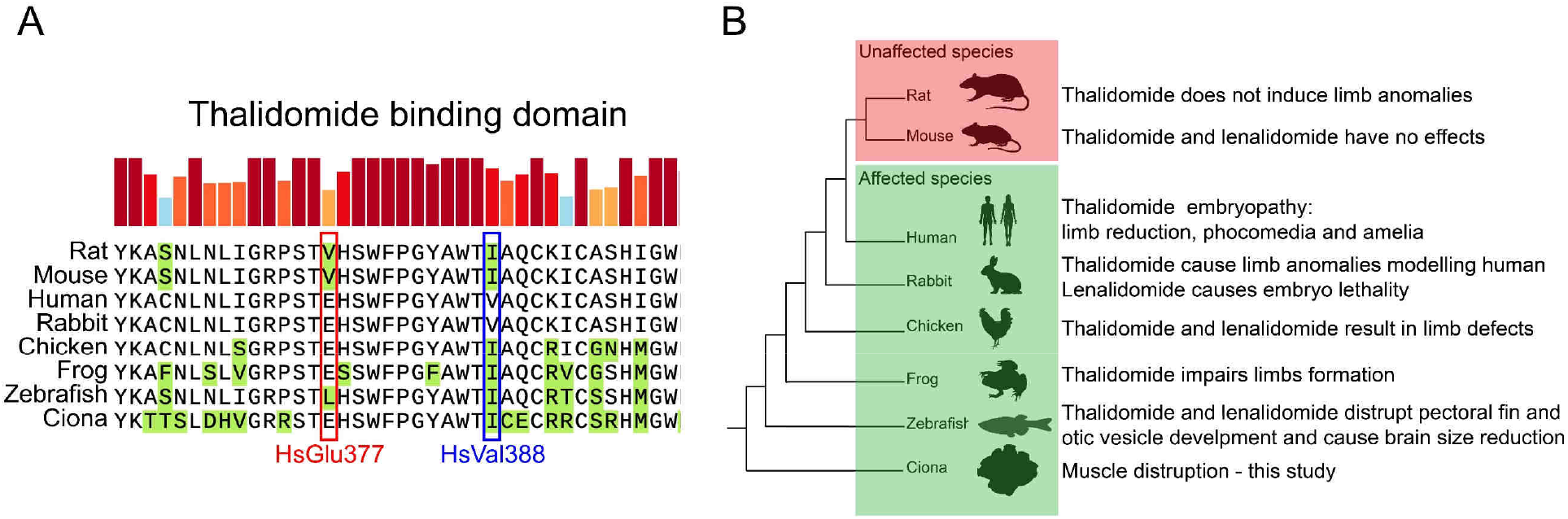
*Ciona Crbn* alignment and phylogenetic tree analysis with other species. **A.** Multiple sequence alignment of the thalidomide binding domain of cereblon among human, rat, mouse, rabbit, chicken, frog, zebrafish and *Ciona*. The red and blue squares correspond to the position of human glutamine 377 (HsGlu377) and human valine 388 (HsVal388) respectively, the two key amino acids for thalidomide and its derivatives interaction with cereblon. Amino acids different from the human sequence are highlighted in green. Both amino acids in the thalidomide resistant species, the mouse and the rat, differ. Conservation scores are plotted on top of the alignment and color-coded (blue: low, red: high). **B.** Phylogenetic tree of the species shown in A. and effects of thalidomide and its analogs exposure are indicated on the left.

**Supp Fig. 2:**
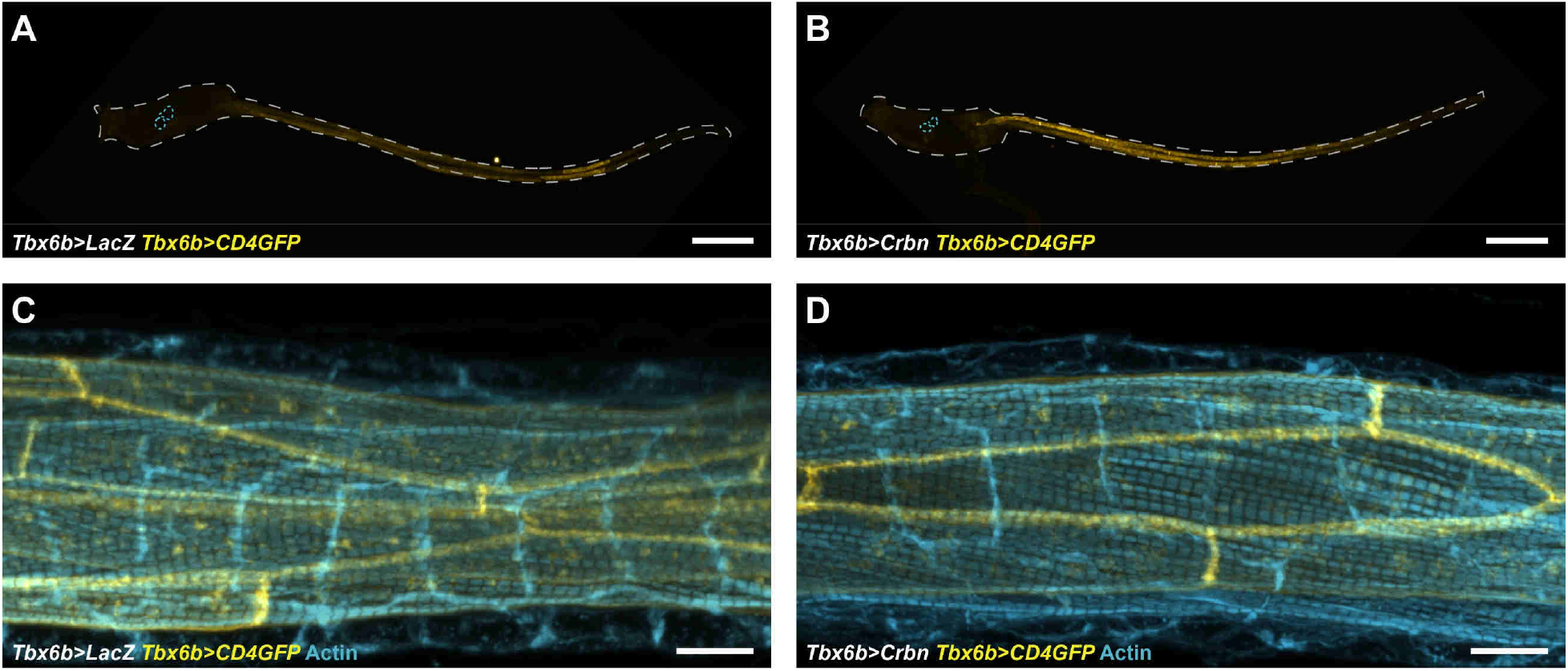
Effect of *Crbn* overexpression in larva muscle cells. **A.** and **B.** Representative larvae expressing either *LacZ* or *Crbn* and the fusion protein CD4GFP (yellow) under the control of muscle-specific *Tbx6b* regulatory sequences. **C.** and **D.** High magnification images showing representative muscle cells stained for actin (blue) in a control larva (*Tbx6b>LacZ*, C.) and in a *Crbn* overexpressing larva (*Tbx6b>Crbn*, D). The muscle membrane is marked by the fusion protein CD4GFP (yellow). Scale bar: A. and B. 100 μm; C. and D. 10 μm.

**Supp Fig. 3:**
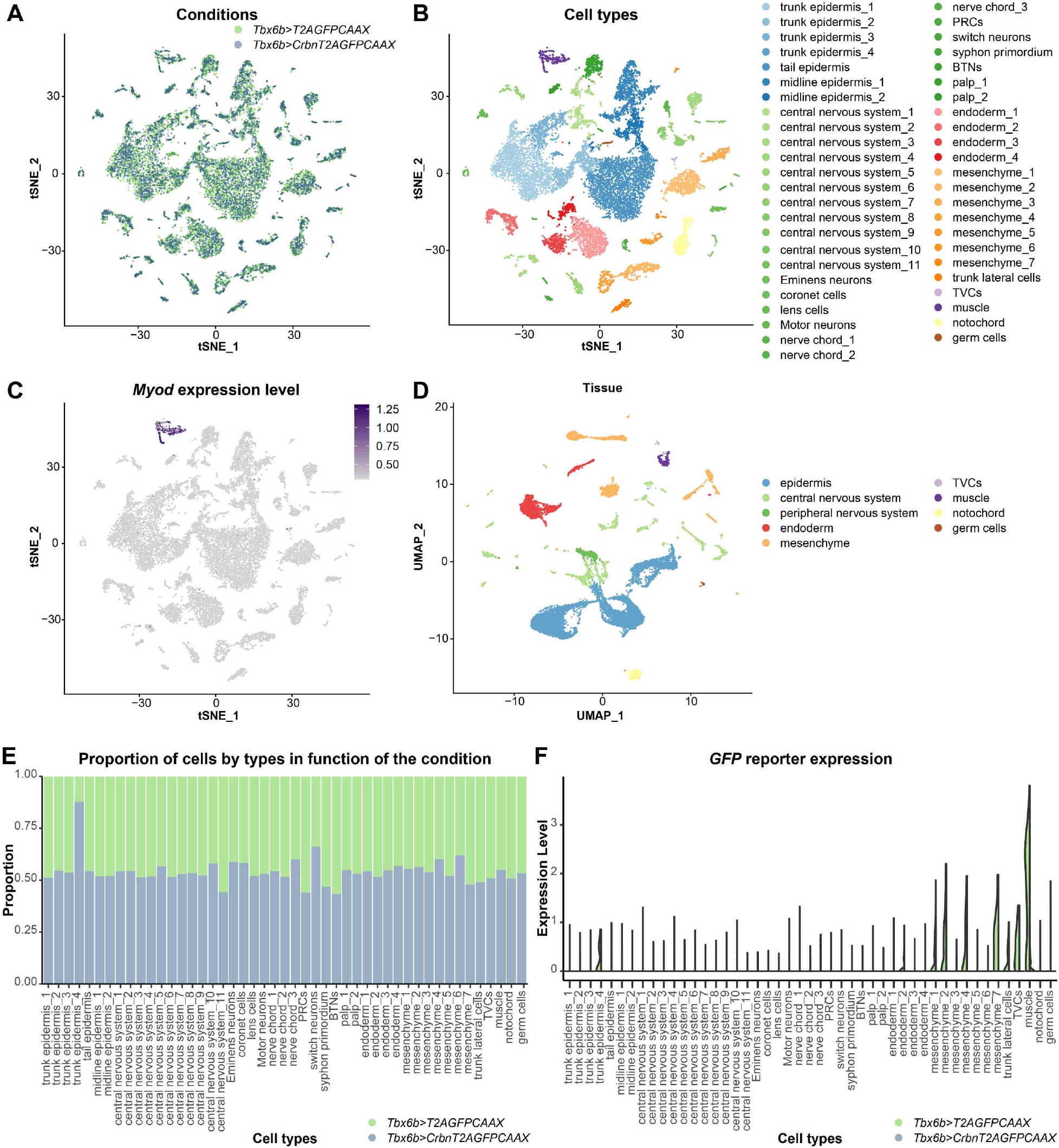
Single cell analysis of *Crbn* overexpressing mid tailbud embryos. **A.** and **B.** t-SNE plots of single cells expressing either *Tbx6b>T2AGFPCAAX* (control condition, n = 9854 cells) or *Tbx6b>CrbnT2AGFPCAAX (Crbn* overexpression condition, n = 8540 cells) at mid tailbud stage. The cells are either color coded based on the condition (A.) or based on their inferred cell types (B.). **C.** t-SNE plot of the same cells showing *Myod* expression level, a marker for the muscle cells. **D.** Uniform manifold approximation and projection (UMAP) co-projection of the same cells color coded by tissue types. **E.** Cell type ratio between control embryos and *Crbn* overexpressing embryos. The number of muscle cells was not affected upon overexpression of *Crbn*. **F.** Violin plot of the expression of the GFP transgene in the different cell types. The following cell types expressed GFP: trunk epidermis_4, mesenchyme_1, mesenchyme_2, mesenchyme_4, mesenchyme_7, TVCs, muscle and were further investigated to analyze the effect of *Crbn* overexpression.

**Supp Fig. 4:**
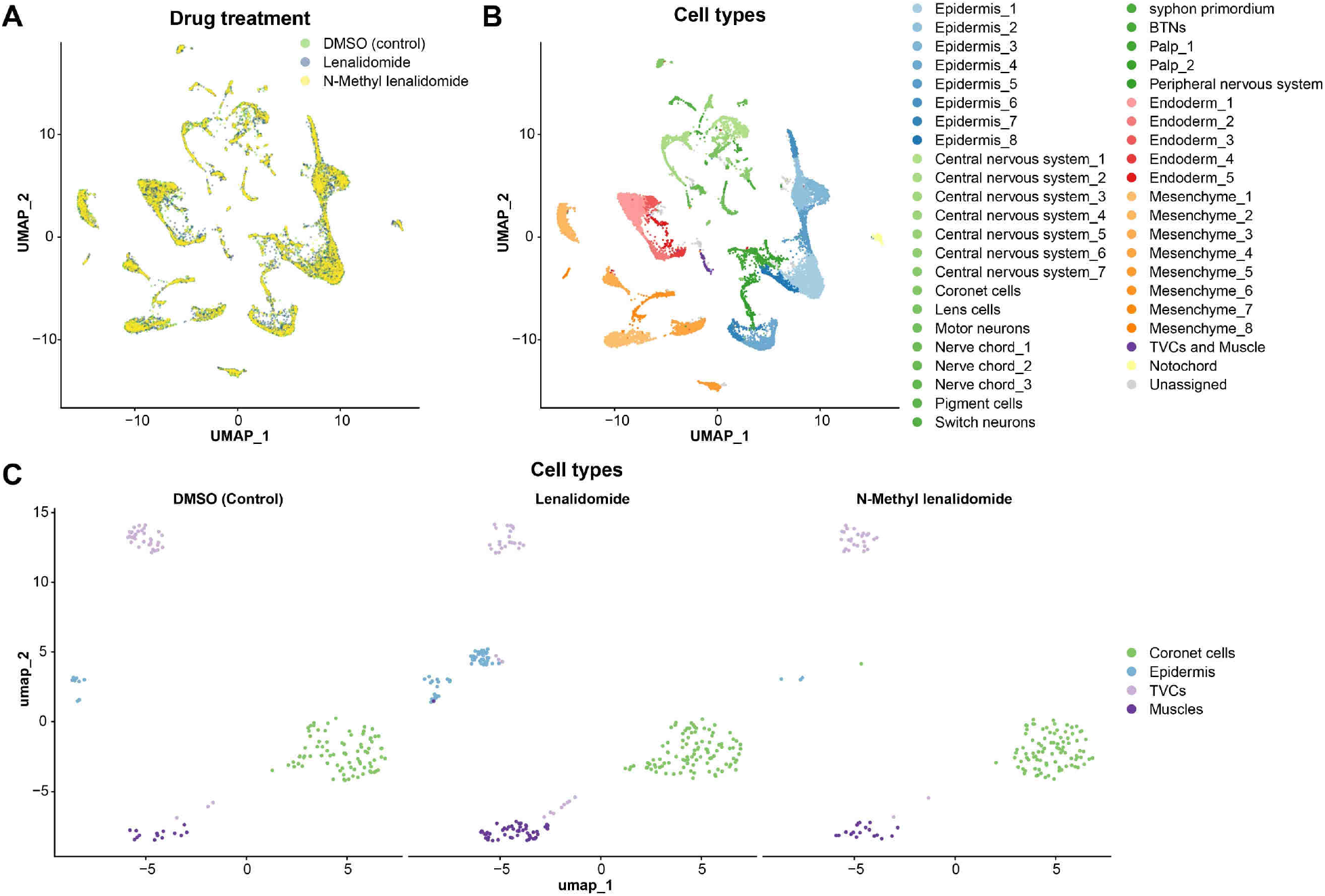
Single cell analysis of late tailbud embryos treated with lenalidomide or N-Methyl lenalidomide. **A.** and **B.** UMAP co-projections of single cells from late tailbud 2 embryos treated either with DMSO (control), 200 μM of lenalidomide or 200 μM of N-Methyl lenalidomide. The cells are color coded by treatment (A.) or by their inferred cell types (B.). **C.** UMAP co-projections of the sub clustering of the coronet cells, TVCs (heart lineage) and muscle cells from the datasets plotted in A. and B. and split by treatment. The cells are color coded by their inferred cell type.

**Supp Fig. 5:**
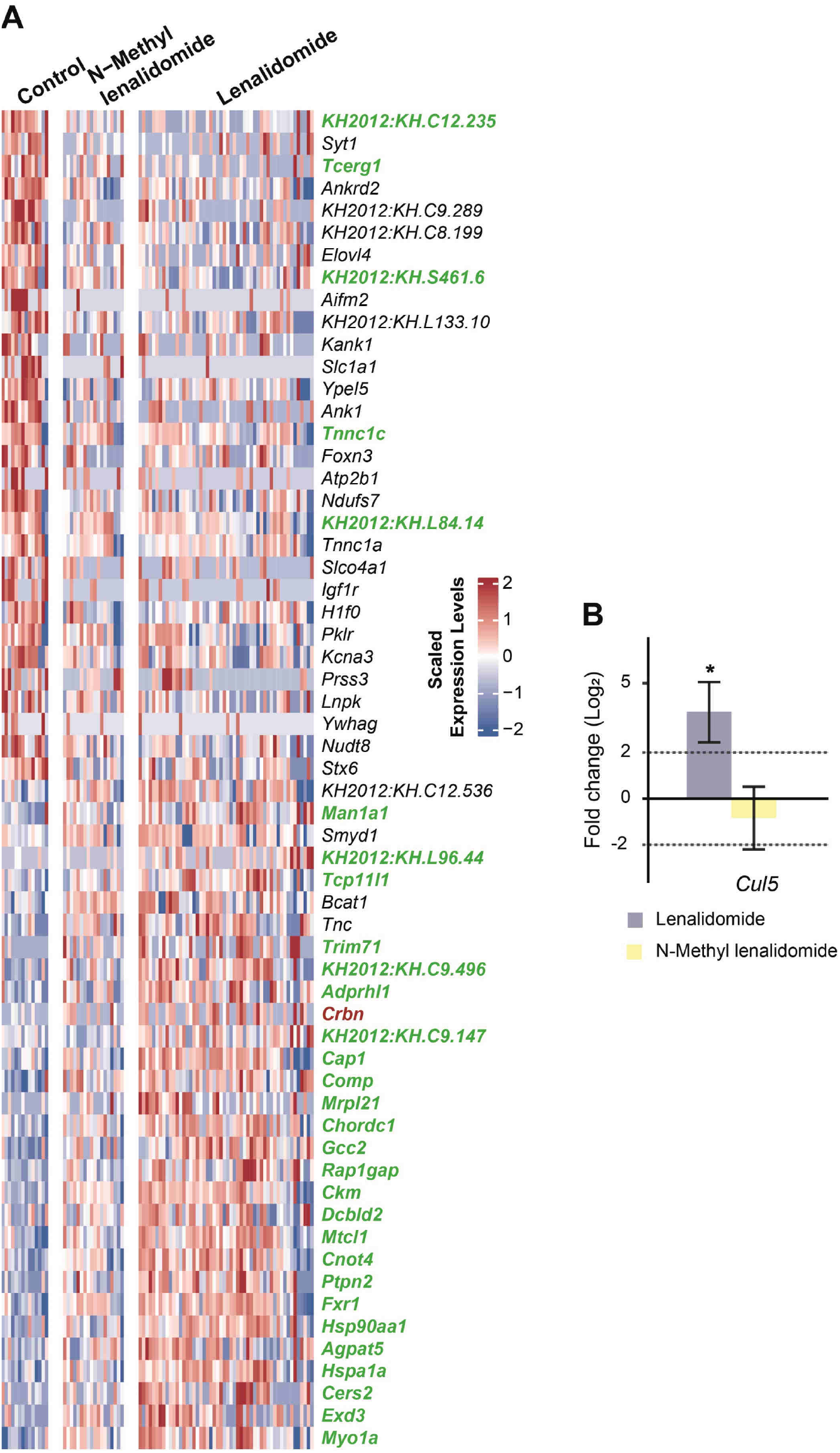
Effect of Lenalidomide and N-Methyl lenalidomide in the muscle cells. **A.** Heatmap of the 30 most differentially expressed genes (up and down) with at least 1.4-fold change and a p value below 0.05 between the muscle cells at the late tailbud 2 stage exposed to 200 μM of lenalidomide and the control muscle cells (DMSO). The expression level in the cells exposed to 200 μM of N-Methyl lenalidomide is also indicated. The genes highlighted in green are also differentially expressed (p value below 0.05) when the lenalidomide exposed muscle cells are compared to the control muscle cells combined to the N-Methyl lenalidomide treated muscle cells. *Crbn* is highlighted in red. **B.** qPCR analysis for *Cul5* on late tailbud 2 embryos treated with either 200 μM of lenalidomide or 200 μM of N-Methyl lenalidomide and compared to DMSO treated embryos (Mean +/- 95% CI: Lenalidomide p value < 2×10^-6^, n = 5; N-Methyl lenalidomide p value = 0.55, n = 5).

